# Screening of candidate host cell membrane proteins involved in SARS-CoV-2 entry

**DOI:** 10.1101/2020.09.09.289488

**Authors:** Norihiro Kotani, Takanari Nakano, Ryusuke Kuwahara

**Affiliations:** Medical Research Center, Saitama Medical University, 38 Morohongo, Moroyama-machi, Iruma-gun, Saitama 350-0495; Department of Biochemistry, Saitama Medical University, 38 Morohongo, Moroyama-machi, Iruma-gun, Saitama 350-0495, Japan; Quantum Wave Microscopy Unit, Okinawa Institute of Science and Technology Graduate University, 1919-1 Tancha, Onna-son, Kunigami-gun, Okinawa, 904-0495, Japan

**Author notes:** To whom correspondence should be addressed: Norihiro Kotani: Medical Research Center, Saitama Medical University, 38 Morohongo, Moroyama-machi, Iruma-gun, Saitama 350-0495, Japan.

**Keywords:** SARS-CoV-2, Membrane protein, Virus entry, SARS-CoV-2 pseudovirus, Proximity proteomics

## Abstract

Coronavirus disease (COVID-19) represents a real threat to the global population, and understanding the biological features of the causative virus, i.e., severe acute respiratory syndrome coronavirus 2 (SARS-CoV-2), is imperative for mitigating this threat. Analyses of proteins such as primary receptors and co-receptors (co-factors), which are involved in the entry of SARS-CoV-2 into host cells, will provide important clues to help control the virus. Here, we identified host cell membrane protein candidates present in proximity to the attachment sites of SARS-CoV-2 spike proteins, using proximity labeling and proteomic analysis. The identified proteins represent key candidate factors that may be required for viral entry. DPP4, Cadherin-17, and CD133 were found to co-localize with cell membrane-bound SARS-CoV-2 spike proteins in Caco-2 cells and thus showed potential as candidate factors. The experimental infection with a SARS-CoV-2 pseudovirus indicated a 2-fold enhanced infectivity in the CD133-ACE2-coexpressing HEK293T cells compared to that in HEK293T cells expressing ACE-2 alone. The information and resources regarding these co-receptor labeling and analysis techniques could be utilized for the development of antiviral agents against SARS-CoV-2 and other emerging viruses.

## INTRODUCTION

Coronavirus disease (COVID-19), caused by severe acute respiratory syndrome coronavirus 2 (SARS-CoV-2), was first reported in Wuhan and now represents a global threat. SARS-CoV-2 is a member of *Coronaviridae*, and many of these coronaviruses have long been known to cause severe respiratory failure (1). Therefore, it is inferred that cells within the respiratory organs are the growth sites for SARS-CoV-2 virions, and analyses of virus receptors within these host cells have been performed. In addition to its ability to infect respiratory organs, SARS-CoV-2 can also infect vascular endothelium and the intestinal tract (2, 3), which indicates the diverse nature of the viral receptors. In SARS-CoV-1 (4, 5) and MERS-CoV (6), certain protease family proteins within host cells have been found to act as primary receptors or co-receptors (co-factors) for these viruses; in particular, ACE2 has been studied as a primary candidate (7). Similarly, SARS-CoV-2 has been reported to target ACE2 and other cell surface proteins (8).

Other membrane proteins involved in virus entry are also of importance in understanding disease pathogenesis. The chemokine receptors CCR5 and CXCR4 have been reported as co-receptors involved in human immunodeficiency virus infection (9–11), and HLA class II receptors function as co-receptors in Epstein-Barr virus infection (12). Researchers working on vaccine and antiviral agent development are interested in the role of viral co-receptors (co-factors), in addition to primary receptors (13). Host cell membrane proteins involved in SARS-CoV-2 attachment and entry can be broadly considered as crucial key factors and therapeutic targets for SARS-CoV-2 infection. In the case of SARS-CoV-2, various host cell membrane proteins other than ACE2 have also been reported as factors that mediate viral entry (14, 15); however, the detailed mechanisms underlying their functions remain unknown. It is speculated that these factors are membrane proteins located in proximity to the viral attachment point (binding site of SARS-CoV-2 spike protein) in the host cell membrane.

The proximity labeling method (16–19) has recently been used to analyze the physiological protein interactions. We developed a simple physiological method, termed <u>E</u>nzyme-<u>M</u>ediated <u>A</u>ctivation of <u>R</u>adical <u>S</u>ource (EMARS) (20), that uses horseradish peroxidase (HRP)-induced radicals derived from arylazide or tyramide compounds (21). The radicals produced through EMARS attack and form covalent bonds with the proteins in proximity to HRP [*e*.*g*., radicals from arylazide: approximately 200–300 nm (20); from tyramide: approx. 20 nm (22)]. The labeled proteins can subsequently be analyzed using an antibody array and/or a typical proteome strategy (23). The EMARS method has been applied for various studies on molecular complexes in the cell membrane (24–31).

Proximity labeling typically analyzes intracellular molecular interactions to provide a measure of the proximity between free proteins. In contrast, EMARS is a tool for analyzing proximity between molecules on the cell surface; this method facilitates the labeling of key factor proteins in proximity to the virus-binding protein on the cell membrane, at the initial stages of infection. Therefore, we speculated that this method would provide a useful tool for identifying key candidate molecules responsible for SARS-CoV-2 infection.

Herein, we identified protein molecules that exist in close proximity to virus spike proteins bound to host cells in the early stage of SARS-CoV-2 infection, using the EMARS method and proteomic analysis. The EMARS reaction was performed in A549 lung cancer cells and Caco-2 cells using the HRP-conjugated recombinant SARS-CoV-2 spike protein (S1-RBD). Caco-2 cells are a commonly used culture cell type that allow for efficient replication of SARS-CoV-2 virions. The labeled proteins identified as candidate membrane proteins were analyzed using proteomics technology. Following infection experiments using a SARS-CoV-2 pseudovirus (pSARS-CoV-2), we generated HEK293T cells coexpressing the identified candidate molecules, together with ACE2, and observed the effects of these candidate molecules on virus infection.

## RESULTS

### Preparation of a SARS-CoV-2 spike protein probe for use in the EMARS reaction

To perform an EMARS reaction, an EMARS probe with HRP conjugated to a given molecule that recognizes a target molecule is required (20). The HRP-conjugated SARS-CoV-2 spike protein corresponds to an EMARS probe in this study (Fig. 1).

**Fig. 1.**
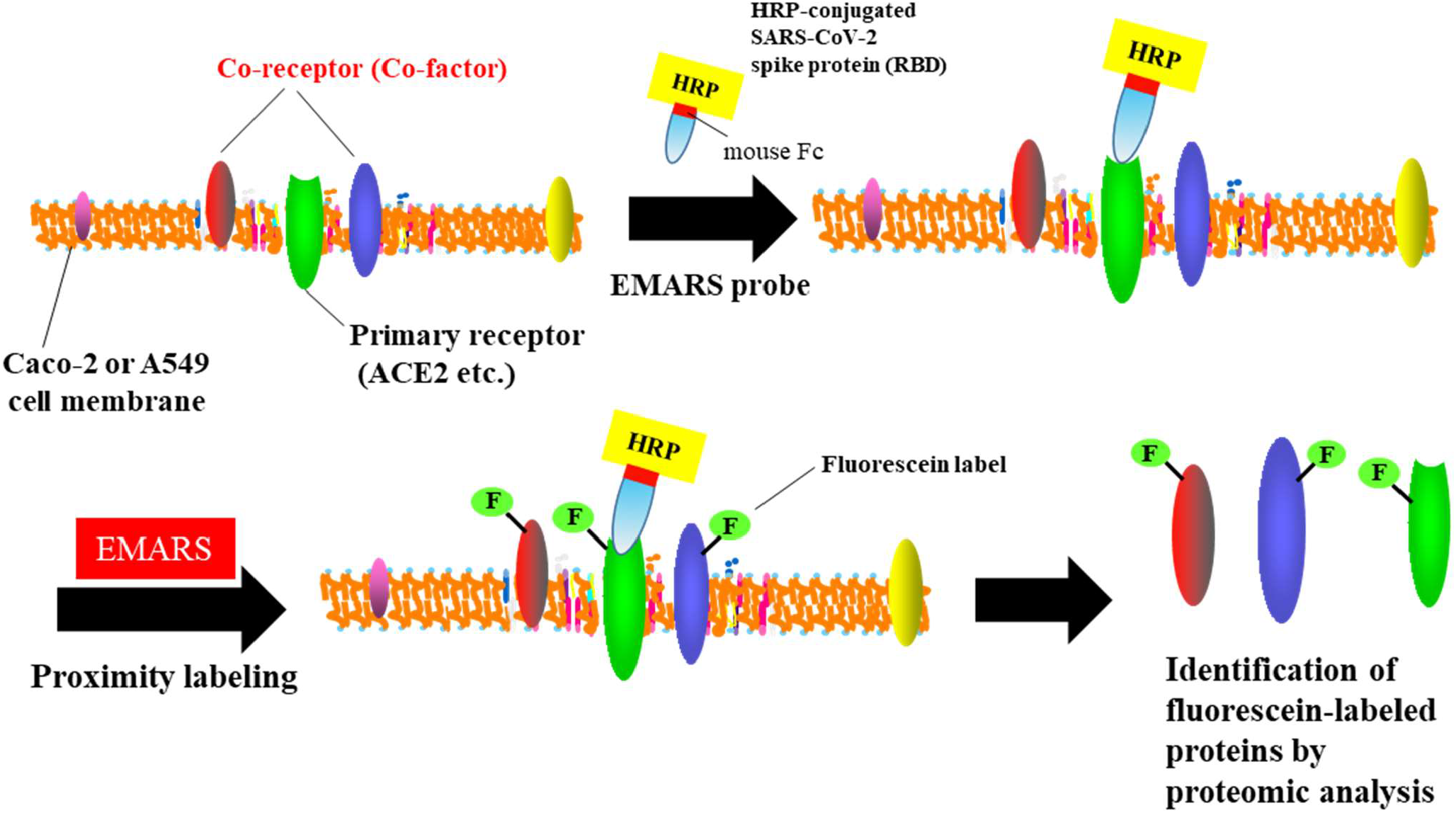
Schematic of the screening for candidate membrane proteins involved in SARS-CoV-2 entry. Schematic illustration of the labeling procedure according to EMARS. After EMARS reaction, the fluorescein-labeled proteins were purified and then analyzed using mass spectrometry.

As ACE2 is listed as the primary receptor for SARS-CoV-2, we examined the expression of ACE2 in Caco-2 and A549 cells. From western blot analysis, a clear band was observed at a molecular weight of approximately 100,000 Da in Caco-2 cells; however, this signal was faint in A549 cells (Fig. 2A). The cells exhibited positive staining for ACE2 antibody, which demonstrated the presence of ACE2 in both cell lines; however, Caco-2 cells exhibited a higher expression of ACE2 than A549 cells (Fig. 2B). Additionally, Caco-2 cells exhibited strong staining of portions of the cell membrane, while A549 cells showed a homogeneous staining pattern (Fig. 2B).

**Fig. 2.**
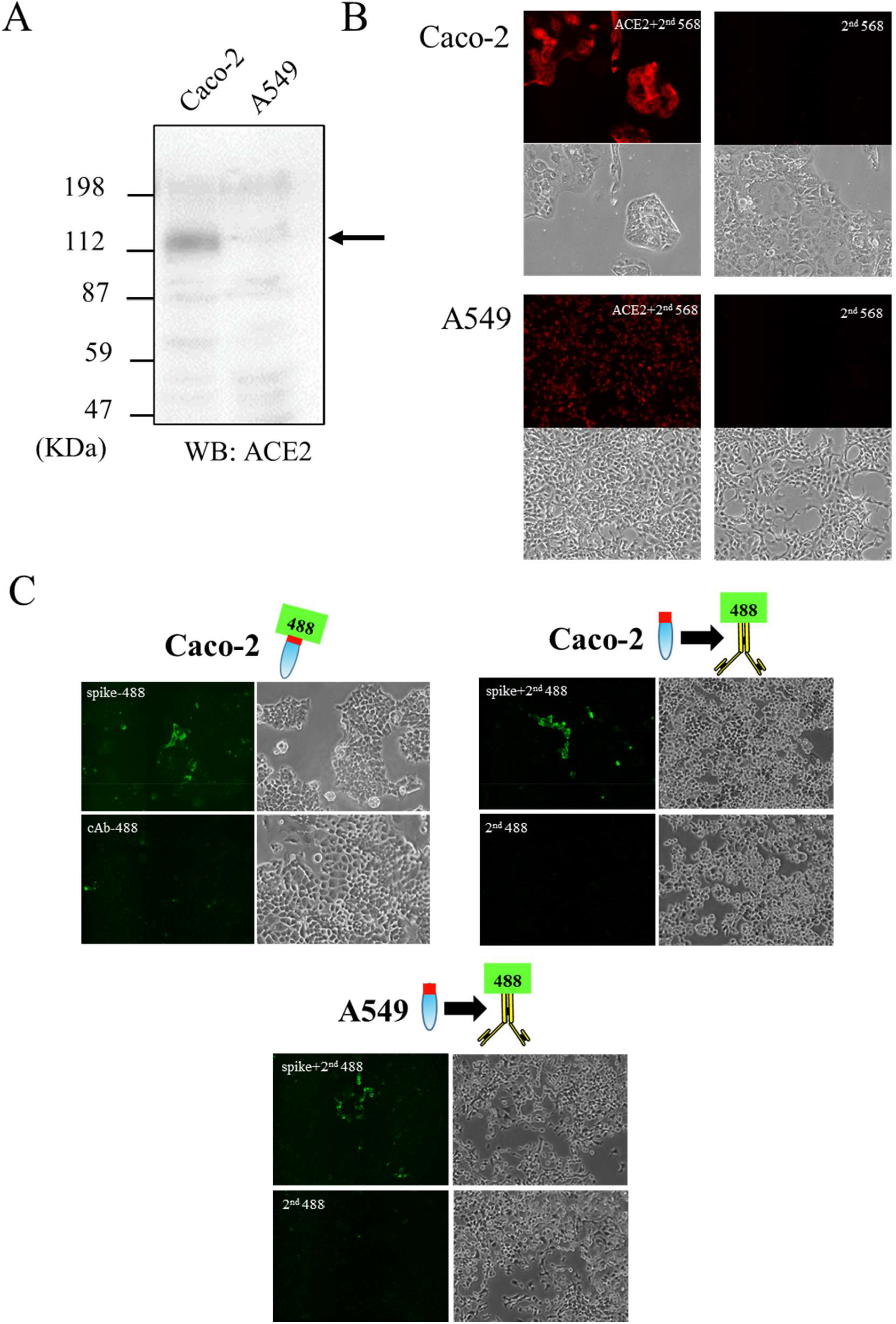
SARS-CoV-2 spike protein-based EMARS probes. (*A*) ACE2 expression in Caco-2 and A549 cells. Western blot analysis of Caco-2 and A549 cell lysates; 10 µg protein samples were subjected to SDS-PAGE (on 10% gels) and stained with anti-ACE2 antibody. Arrows indicate bands of the ACE2 protein. (*B*) Immunocytochemical staining of ACE2 in Caco-2 and A549 cells. Staining with the anti-ACE2 antibody (ACE2+2^nd^ 568) was performed as described in *Experimental procedure*. Negative control samples (2^nd^ 568) were also prepared simultaneously. (*C*) Immunocytochemical staining of SARS-CoV-2 spike proteins in Caco-2 and A549 cells. Staining of monovalent Alexa Fluor 488-labeled spike proteins (spike-488) and the two-step staining (spike protein followed by Alexa Fluor 488 secondary antibody; spike+2^nd^ 488) were performed with DIC images. Negative control samples (cAb-488 or 2^nd^ 488) were also prepared simultaneously.

The spike protein used in this study was a receptor-binding domain (RBD) that is present in the S1 protein of SARS-CoV-2 and possesses a mouse immunoglobulin Fc region (mFc) at the C-terminus (total 457 a. a.). We used the mFc sequence to conjugate HRP. We initially examined whether the spike protein binds to membrane receptors on Caco-2 and A549 cells. Caco-2 cells were treated with Alexa Fluor 488 fluorescent dye-conjugated monovalent spike proteins created using a commercial labeling kit, and labeling was then observed using a fluorescent microscope. We simultaneously prepared the sample that was created through incubation with intact spike protein-mFc, followed by incubation with anti-mouse IgG-Alexa Fluor 488. Both samples exhibited binding of spike protein-mFc to the cells; however, the latter two-step staining procedure was found to be more effective (Fig. 2C). Based on these results, a method for EMARS in this study was adopted wherein the SARS-CoV-2 spike protein was directly applied to living cells and then treated with HRP-conjugated anti-mouse IgG. Similar results were obtained using A549 cells; however, binding of the SARS-CoV-2 spike protein was weaker than that observed in Caco-2 cells (Fig. 2C).

### EMARS and proteomic analysis revealed numerous candidate proteins

The EMARS reaction was performed in these cells using the EMARS probe described above. Following the EMARS reaction, the labeled molecules were subjected to SDS-PAGE. The gels used for electrophoresis can be analyzed directly using a fluorescence image analyzer when the labeled molecules are present in large amounts; however, the bands for the EMARS sample prepared using the SARS-CoV-2 spike protein could not be clearly detected. The labeled molecules were therefore observed with western blot analysis using the anti-fluorescein antibody. The EMARS reaction was first performed in Caco-2 cells using HRP-conjugated cholera toxin subunit B as the positive control, as this conjugate is known to bind to a lipid raft structure on the cell membrane and can subsequently label numerous cell surface proteins (Fig. 3A). We then performed the EMARS reaction, using the SARS-CoV-2 spike protein, in addition to negative control experiments. Although weak bands were observed in the negative control (treated with anti-mouse IgG-HRP antibody alone; hereinafter referred to as spike [-] sample), the combination of both SARS-CoV-2 spike protein and anti-mouse IgG-HRP antibody (hereinafter referred to as spike [+] sample) yielded significant bands (Fig. 3A). As the signal was weak, compared to that obtained using the cholera toxin probe, it was possible that certain specific cell membrane proteins were labeled in a limited manner in the samples prepared using the SARS-CoV-2 spike protein probe. For A549 cells, a number of labeled proteins were detected in the spike [+] sample; however, the band pattern was different from that obtained from Caco-2 cells (Fig. 3B).

**Fig. 3.**
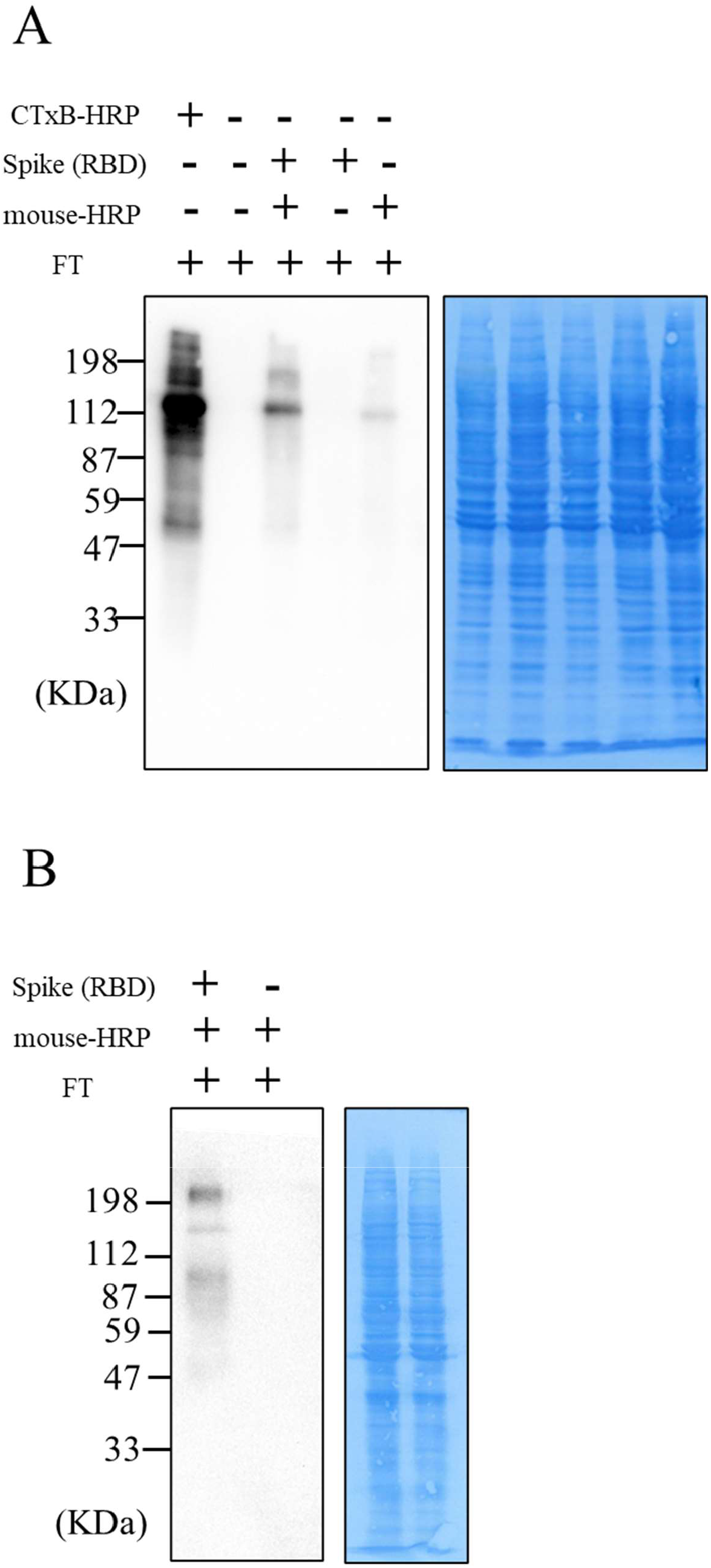
Proximity labeling near the cell membrane-bound SARS-CoV-2 spike protein. (*A, B*) Fluorescein-labeled proximal proteins around cell membrane-bound SARS-CoV-2 spike proteins. The EMARS reaction described in the “Experimental procedure” was performed in Caco-2 (*A*) and A549 (*B*) cells using a spike protein (*Spike (RBD)*) and HRP-conjugated anti-mouse IgG (*mouse HRP*). The EMARS products were subsequently subjected to Western blot analysis to detect fluorescein-labeled proteins as candidate proximal proteins. In Caco-2 cells, HRP-conjugated Cholera Toxin B Subunit B (*CTxB-HRP*) was used for EMARS reaction as the positive control for membrane protein labeling. For loading controls, the PVDF membrane was stained with Coomassie Brilliant Blue after western blot analysis (right column)

These labeled proteins were subjected to proteomic analysis. There was a possibility that identification may not be sufficiently achieved using mass spectrometry, as the number of labeled molecules was likely to be lesser than that obtained using the CTxB-HRP probe. Therefore, multiple independent experiments were performed (twice for Caco-2 cells and three times for A549 cells), and the samples were combined and used for analysis. Moreover, this experiment was performed in duplicate.

Spike [+] samples and spike [-] samples (used as the negative control) were both prepared from each cell line. For shotgun analysis using mass spectrometry, the labeled molecules were purified via immunoprecipitation with an anti-fluorescein antibody. A total of 181 (first MS analysis) and 315 (second MS analysis) proteins were detected in the spike [+] samples from Caco-2 cells, whereas 59 and 230 proteins were detected from the spike [-] samples (Supporting tables 1 and 2). A total of 184 and 263 proteins were detected in the spike [+] samples from A549 cells, and 186 and 150 proteins were detected from the spike [-] samples (Supporting tables 3 and 4). The molecules detected in the spike [-] sample may be proteins that were non-specifically adsorbed during the purification process. In particular, for unknown reasons, some spike [-] samples contained many suspected non-specific binding proteins. In this study, membrane proteins that were present in the spike [+] sample but not in the spike [-] sample, for each cell line, were preferentially categorized as the most likely candidates in this study. In Caco-2 cells, 65 types of cell surface membrane proteins, including the known SARS-CoV-2 host factors ACE2, DPP4, integrin, and CEACAM, were identified using combined data from duplicate experiments (Supporting table 5). Among these candidates, we listed other less implicated proteins in SARS-CoV-2 infection, such as Cadherin 17 (Table 1). In A549 cells, 18 types of membrane proteins, including known SARS-CoV-2 host factors (Supporting table 5), and less reported proteins, such as Contactin-1 (Table 2), were identified.

### The identified membrane proteins co-localized with the SARS-CoV-2 spike protein

We next examined whether the identified membrane proteins co-localized with the SARS-CoV-2 spike protein bound to Caco-2 cell membrane. Caco-2 cells were treated with SARS-CoV-2 spike proteins, followed by staining with antibodies against ACE2, CD133, Cadherin 17, and DPP4 candidates. Representative images of ACE2, CD133, Cadherin 17, and DPP4 are shown in Fig. 4. It was found that all these proteins (red signals) were at least expressed in Caco-2 cells and co-localized with the SARS-CoV-2 spike protein (green signals). Interestingly, despite the observation that these proteins were expressed abundantly, the co-localized area was limited to specific membrane sites.

**Fig. 4.**
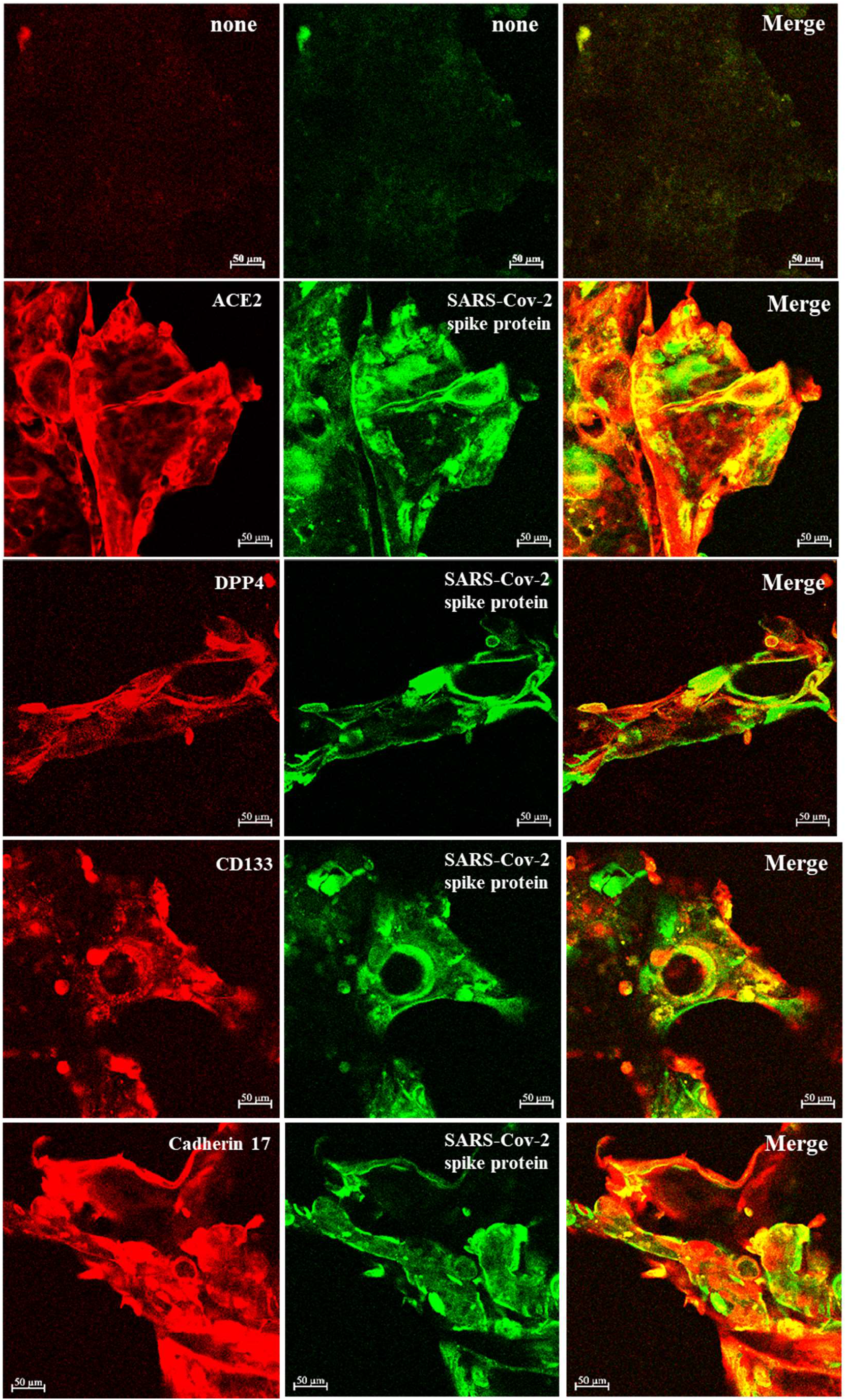
Co-localization of the identified proteins with cell membrane-bound SARS-CoV-2 spike proteins. Representative images of co-localization with SARS-CoV-2 spike proteins and the identified membrane proteins. Caco-2 cells were co-stained for SARS-CoV-2 spike protein (green; middle column) and the antibodies recognizing ACE2, CD133, Cadherin 17, and DPP4 (Red; left column). Then, the resulting specimens were stained with appropriate secondary antibodies and subsequently observed using confocal microscopy (20× objective). Co-localization is indicated in yellow in the “Merge” images (right column).

These colocalizations were subsequently examined via transmission electron microscopy (TEM). DPP4, CD133, CDH17, and VAPA (instead of ACE2) were labeled with 10 nm gold particles (yellow arrowhead), and the spike protein was labelled with 20 nm gold particles (red arrow). DPP4, CD133, CDH17, and VAPA localized close to the binding site of the spike proteins, and this finding was consistent with the results of confocal microscopy analysis (Fig. 5).

**Fig. 5.**
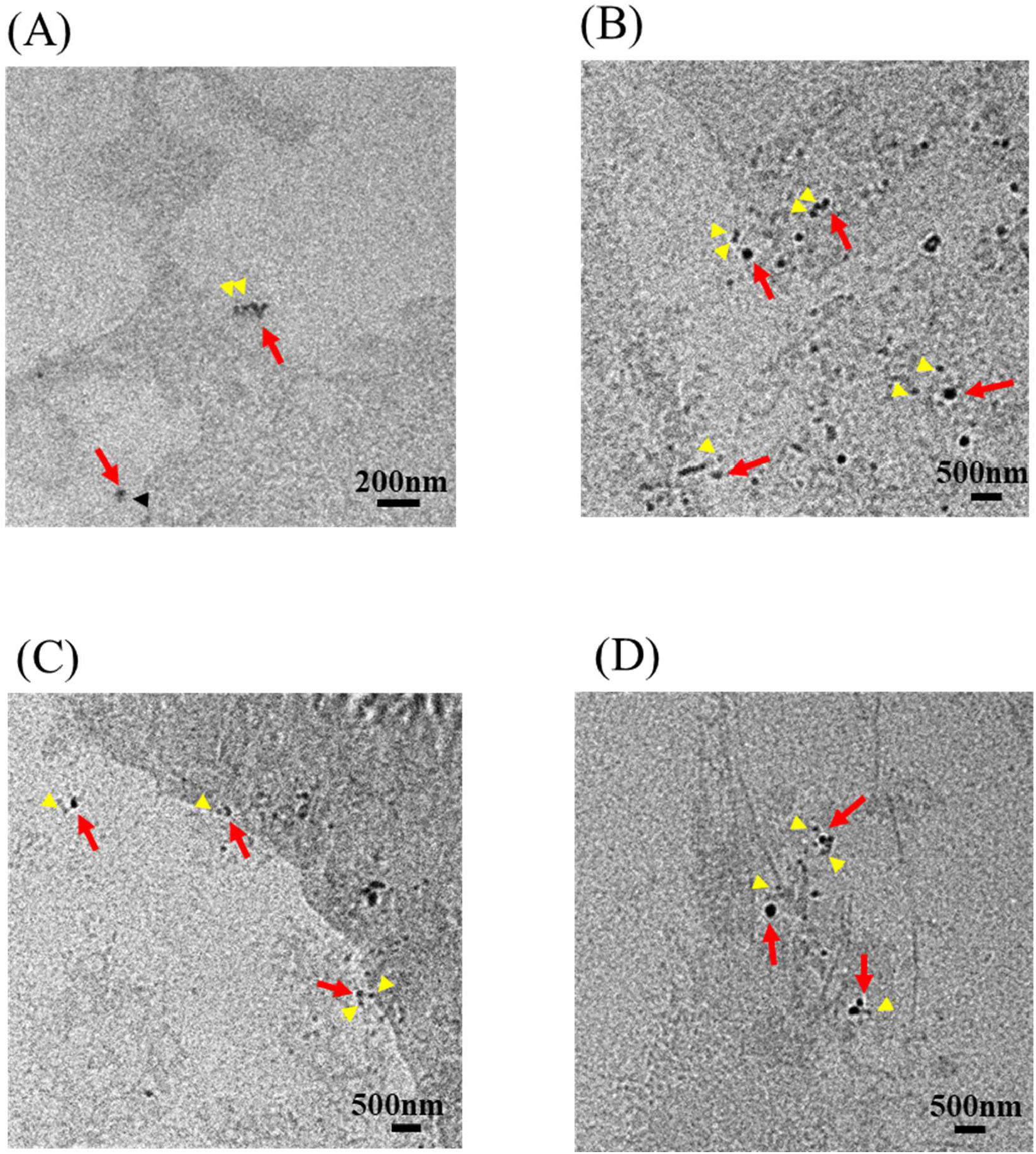
Candidate proteins located near SARS-CoV-2 spike proteins. (*A to D*) Morphological observation of SARS-CoV-2 spike proteins and the identified membrane proteins. Caco-2 cells observed using electron microscopy. Cultured Caco-2 cells were fixed and co-stained with the SARS-CoV-2 spike protein (indicated as 20 nm particles), and candidate molecules identified. CD133 (*A*), DPP4 (*B*), CDH17 (*C*), and VAPA (*D*) are indicated as 10 nm particles. Red arrows indicate the locations of SARS-CoV-2 spike proteins. Yellow arrow heads indicate the location of each candidate protein. Scale bar; 200 or 500 nm.

### Preparation of HEK293T transfectant cells expressing ACE2 and/or candidate proteins

To elucidate whether the candidate proteins affect the efficacy of SARS-CoV-2 infection, HEK293T cells expressing ACE2 and/or candidate proteins were prepared for the infection assay of SARS-CoV-2. We first prepared two types of ACE2-expressing HEK293T cells using a PCMV3 expression vector system (P-ACE2; Supporting Fig. 1A) or a lentivirus expression system (L-ACE2; Supporting Fig. 1B). Flow cytometric analysis revealed sufficient expression levels of ACE2 in both cell lines (Supporting Fig. 1C). As ACE2 is a primary receptor for SARS-CoV-2, we confirmed the binding capacity of the SARS-CoV-2 spike protein, and strong binding was observed in both cell lines (Supporting Fig. 1D). The binding amount was slightly higher in L-ACE2 (Supporting Fig. 1E), demonstrating a correlation with ACE2 expression.

Furthermore, we prepared single transfectant HEK293T cells expressing candidate proteins (CD133, CDH17, and VAPA), using a lentivirus expression system (Supporting Fig. 2). Binding of the SARS-CoV-2 spike protein was not detected in these cells (Supporting Fig. 2), indicating that these candidate proteins are not the primary receptors of SARS-CoV-2.

To perform cotransfection of CD133, CDH17, and VAPA with ACE2-expressing cells, the lentivirus system used in the above-mentioned experiment was subsequently applied to P-ACE2 cells (Supporting Fig. 3A). Glypican-3 (GPC3), which was detected in both the spike [+] and spike [-] samples in MS analysis, was also cotransfected in P-ACE2 cells as the negative control (Supporting Fig. 3A).

CD133-, CDH17-, and VAPA-coexpressing P-ACE2 cells (P-ACE2-CD133, -CDH17, and -VAPA) were subjected to the binding assay of the SARS-CoV-2 spike protein (Supporting Fig. 3B). The extent of spike protein binding to an ACE2 molecule was higher than that to P-ACE2 cells; however, no statistically significant difference was observed (Supporting Fig. 3C).

### Candidate membrane proteins partially affect the infection of SARS-CoV-2 pseudovirus

A lentivirus-based pSARS-CoV-2 was generated to perform the infection assay in each cell (Fig. 6A). Since this pSARS-CoV-2 was capable of expressing GFP in infected cells (Fig. 6A), the infected cells could be determined by assessing the number of GFP-positive cells.

**Fig. 6.**
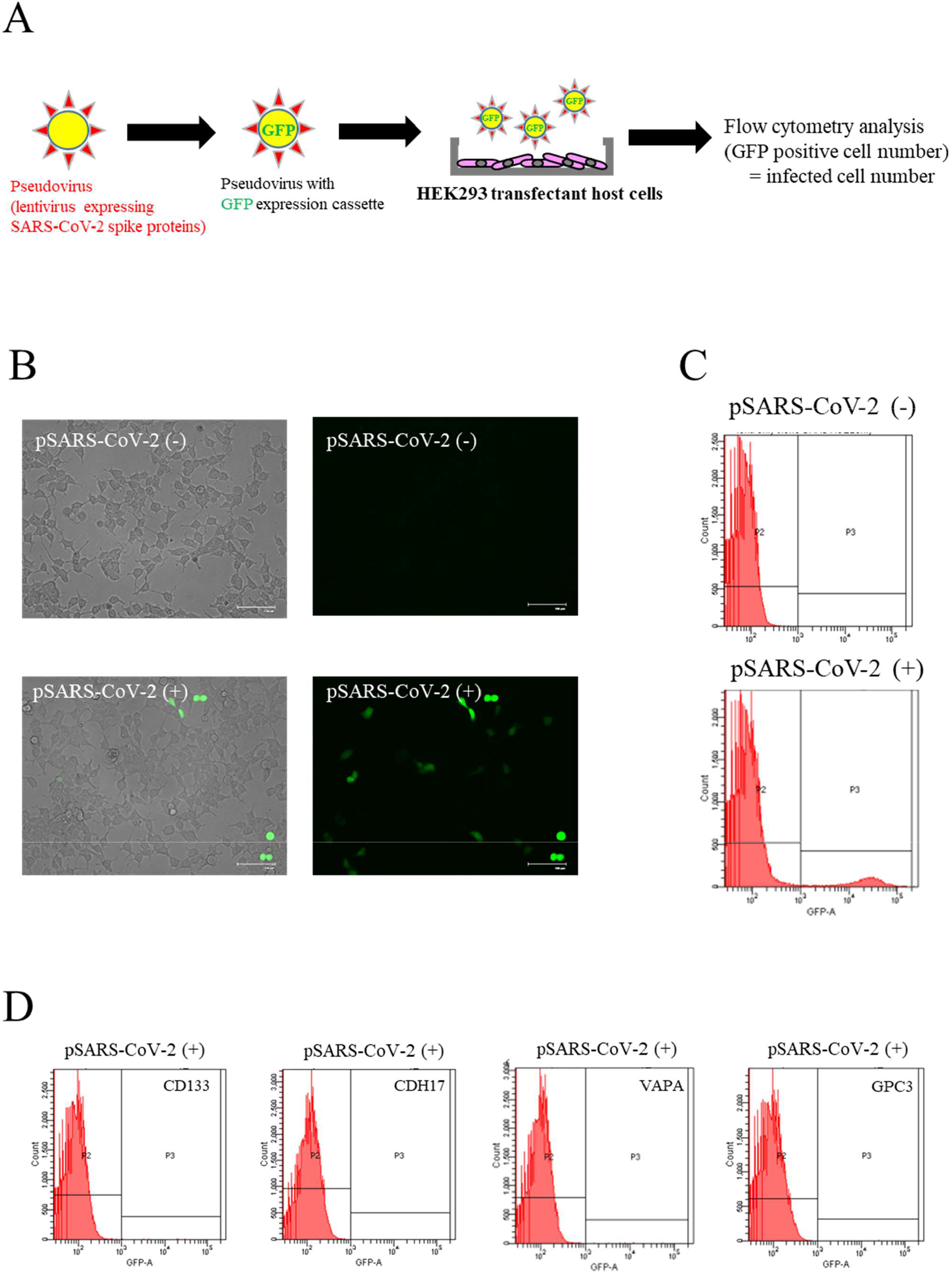
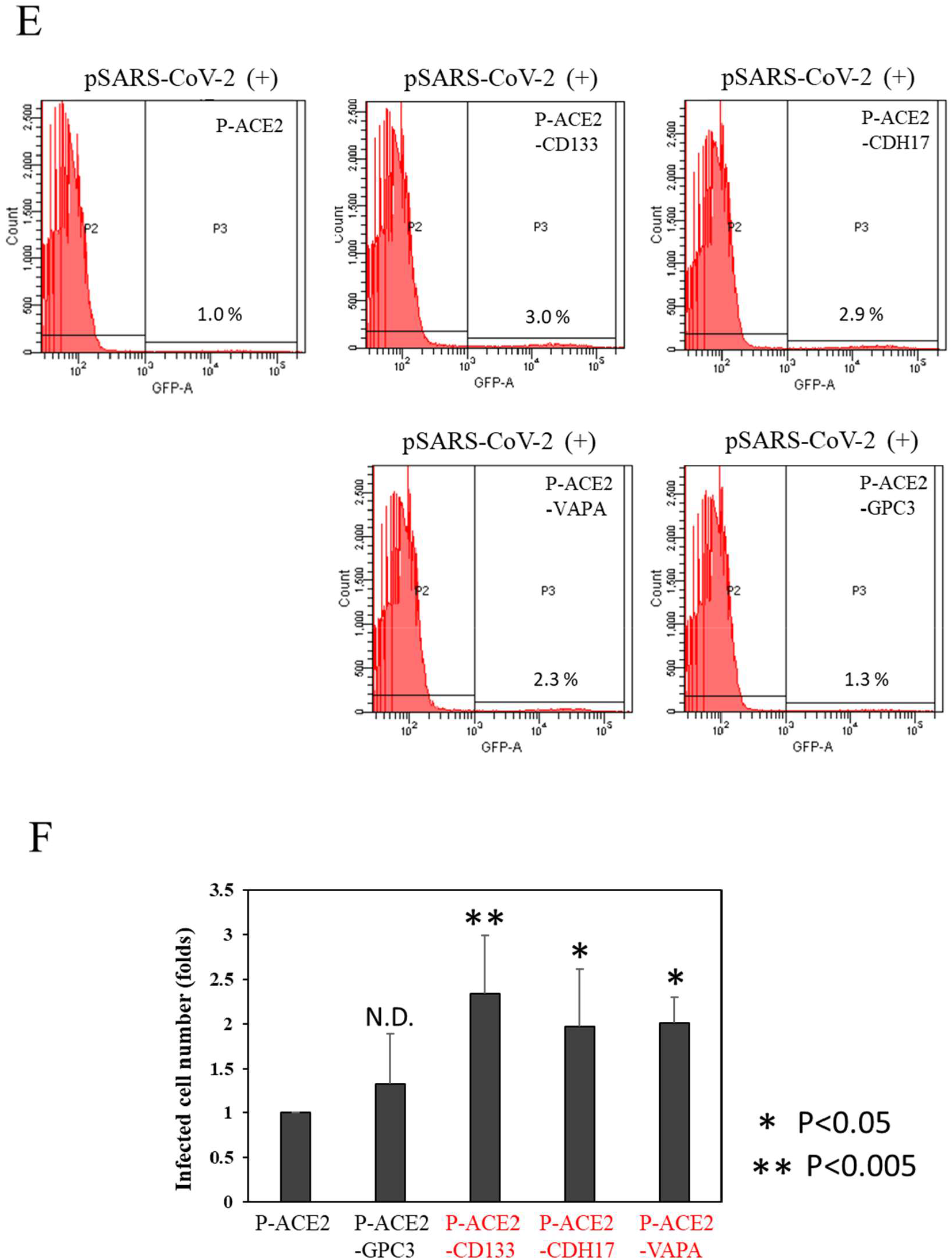
*In vitro* infection assay of SARS-CoV-2 pseudovirus. (*A*) Schematic illustration of the assay procedure using HEK293T transfectant host cells. (*B*) Representative images of GFP-positive P-ACE2 cells after pSARS-CoV-2 infection. ACE2-expressing HEK293T cells were treated (pSARS-CoV-2 (+)) or not treated (pSARS-CoV-2 (-)) with pSARS-CoV-2, followed by fluorescein microscopic observation. Two independent experiments were carried out. (*C-E*) Flow cytometric analysis of pSARS-CoV-2-infected cells. P-ACE2 cells (*C*), candidate protein-single expressing cells (*D*), and candidate protein-coexpressing P-ACE2 cells (*E*) were analyzed using BD FACS Canto II. GFP-positive cells were defined as the infected cells with a GFP fluorescence intensity of 10^3^ or higher (P3 area). Two (*D*) or five (*C* and *E*) independent replications were carried out in each experiment. (*F*) Increase in pSARS-CoV-2 infection in candidate protein-coexpressing P-ACE2 cells. The number of GFP-positive cells in each cell was quantified using flow cytometry. The number of infected cells (GFP-positive) in P-ACE2– CD133, –CDH17, and –VAPA was significantly higher than that in P-ACE2 cells (*P < 0*.*05* or *P* < 0.005; Dunnett’s test), but not in P-ACE2-GPC3 (N.D.) as the negative control.

The treated cells were observed, using fluorescence microscopy, 60 h after the addition of pSARS-CoV-2. Some GFP-positive cells were observed in pSARS-CoV-2-treated wells (Fig. 6B). For the quantification of GFP-positive cells, the treated cells were subsequently subjected to flow cytometric analysis (Fig. 6C). An independent experiment revealed that in the infection assay, GFP-positive cells accounted for about 1 to 5% of the total cells. In contrast to this, when single transfectant cells expressing the candidate proteins CD133, CDH17, and VAPA were applied to the analysis, almost no GFP-positive cells were observed (Fig. 6D).

The same infection assay revealed that the P-ACE2 cells coexpressing CD133, CDH17, and VAPA had approximately 2 to 3-fold significantly more number of GFP-positive cells compared to the P-ACE2 cells, but not the P-ACE2-GPC3 negative control cells, indicating the ability of these candidate proteins to enhance pSARS-CoV-2 infection (Fig. 6E and 6F).

## DISCUSSION

Proteomic-based analysis has opened up the possibility for the use of host cell proteins involved in SARS-CoV-2 replication as therapeutic targets (32). However, as viral infection initially occurs through the host cell membrane, it is also imperative to focus on membrane proteins, such as the primary viral receptor. In particular, previous studies on viruses have suggested that lipid rafts, which are assembly structures consisting of several membrane proteins, are important for viral entry into host cells (33). Moreover, some studies have demonstrated that the EMARS reaction used in this study is effective for identifying proximity proteins, including the membrane proteins in the lipid raft structure (34). It was therefore speculated that EMARS is suitable for investigating crucial membrane proteins located in the proximity region of the cell membrane-bound SARS-CoV-2 spike protein involved in virus entry.

It has been reported that SARS-CoV-2 expands more easily in Caco-2 cells than in A549 cells (8, 35). These differences could be due to the variation in ACE2 expression levels in these cells. In the current study, ACE2 expression was more clearly observed in Caco-2 cells than in A549 cells (Fig. 2A and 2B), thus supporting the binding capacity of the SARS-CoV-2 spike protein in this study. In contrast, it has also been reported that the expression levels of ACE2 in A549 cells are relatively high (36), and that virus attachment could occur even if the replication ability was low (35). Regardless, as the binding of the spike protein to A549 cells was observed in this study, it was concluded that the experiment could be performed in both Caco-2 and A549 cells.

For preparation of the EMARS probe, the binding capacity of the SARS-CoV-2 spike protein toward the cell membrane was assayed. We expected to observe clear spike protein staining in Caco-2 cells owing to the high ACE2 expression in this cell line. However, only a portion of the cell population could bind to the SARS-CoV-2 spike protein, whereas the other cells did not exhibit clear binding (Fig. 2C). Previous studies examining SARS-CoV-2 infection observed both infected and non-infected cells even in homogeneous cell lines (2, 35). In HeLa cells that overexpress ACE2, not all cells were infected (37). Moreover, the cytological images of spike protein used in this study were similar to those of SARS-CoV-2 infection reported by other studies. The distinction of the binding among cultured cells is presumed to be due to the condition of each cell population (*e*.*g*., the cell cycle and expression levels of co-receptor [co-factors] membrane proteins); however, the actual cause remains unknown.

The staining experiments implied that the amount of membrane proteins labeled by the EMARS reaction may be small. The labeling of molecules by EMARS was confirmed in both Caco-2 cells and A549 cells (Fig. 3A and 3B); however, the number of labeled molecules was less than expected and could only be detected using the Western blot analysis. Considering the loss of the labeled protein in the purification process and the sensitivity of shotgun analysis via mass spectrometry, it was necessary to combine multiple samples obtained from multiple independent experiments despite the non-specific adsorption that occurs during the purification process. In A549 cells, a number of proteins that were expected to be non-specific adsorbed proteins were detected in the negative control sample and were not subjected to the EMARS reaction (spike [-]). We therefore decided to select the candidate membrane proteins using a subtraction method that incorporated the membrane proteins present in the EMARS reaction samples (spike [+]) in both cells (Supporting Table 1 to 5). Although there was a difference in the expression of candidate proteins between Caco-2 cells and A549 cells, it was speculated that the membrane proteins that act as co-factors for SARS-CoV-2 entry were dependent on cell origin. As SARS-CoV-2 is known to infect multiple cell types (2, 3, 35), these differences appear to be events that must be considered in future SARS-CoV-2 research. In contrast, some identical proteins between Caco-2 and A549 cells, including the transferrin receptor protein and others, were detected. These are ubiquitous proteins; however, they may still play an important role by acting as a common target against SARS-CoV-2 infection. It should also be noted that ubiquitous and highly expressed cell membrane proteins are more likely to be detected in proteomic analysis. Another factor is that ACE2, which is the primary receptor of SARS-CoV-2, was detected in Caco-2 cell as a candidate protein in this study (Supporting Table 5), indicating that our strategy can capture molecules in proximity to the spike protein. To narrow down the candidate proteins, the molecules that have never been implicated in SARS-CoV-2 infection were listed (Tables 1 and 2).

**Table 1.** Selected candidates for proximal membrane proteins around the SARS-CoV-2 spike protein in Caco-2 cell surface.

| Accession No. | Protein Name | Score* | Peptide |
| --- | --- | --- | --- |
| Q12864 | Cadherin-17 | 422 | 11 |
| O43490 | Prominin-1 (CD133) | 274 | 7 |
| Q9HBB8 | Cadherin-related family member 5 | 273 | 4 |
| Q5ZPR3 | CD276 antigen | 150 | 4 |
| P21796 | Voltage-dependent anion-selective channel protein 1 | 145 | 4 |
| Q9BYE9 | Cadherin-related family member 2 | 109 | 4 |
| Q9NZU0 | Leucine-rich repeat transmembrane protein FLRT3 | 105 | 5 |
| P13987 | CD59 glycoprotein | 98 | 2 |
| P50895 | Basal cell adhesion molecule | 88 | 2 |
| O00592 | Podocalyxin | 73 | 3 |
| P05026 | Sodium/potassium-transporting ATPase subunit beta-1 | 72 | 2 |
| P15328 | Folate receptor alpha | 71 | 1 |
| Q9BY67 | Cell adhesion molecule 1 | 70 | 2 |
| Q86SQ4 | Adhesion G-protein coupled receptor G6 | 58 | 2 |
| P78310 | Coxsackievirus and adenovirus receptor | 56 | 1 |
| P04156 | Major prion protein | 55 | 1 |
| Q8WW52 | Protein FAM151A | 53 | 1 |
| Q9Y3Q0 | N-acetylated-alpha-linked acidic dipeptidase 2 | 51 | 1 |
| P08174 | Complement decay-accelerating factor | 48 | 2 |
| Q53RT3 | Retroviral-like aspartic protease 1 | 48 | 1 |
| Q9Y277 | Voltage-dependent anion-selective channel protein 3 | 48 | 1 |
| O75915 | PRA1 family protein 3 | 44 | 1 |
| P16444 | Dipeptidase 1 | 41 | 1 |
| P08195 | 4F2 cell-surface antigen heavy chain | 38 | 2 |
| P0C7N4 | Transmembrane protein 191B | 38 | 1 |
| Q16651 | Prostasin | 35 | 1 |
| Q8NFZ8 | Cell adhesion molecule 4 | 32 | 1 |
| P48960 | Adhesion G protein-coupled receptor E5 | 27 | 1 |
| Q9P0L0 | Vesicle-associated membrane protein-associated protein A | 26 | 1 |
| Q08174 | Protocadherin-1 | 25 | 1 |
| Q14118 | Dystroglycan 1 | 25 | 1 |
| P05023 | Sodium/potassium-transporting ATPase subunit alpha-1 | 23 | 1 |
| Q13641 | Trophoblast glycoprotein | 22 | 1 |
| P13796 | Plastin-2 | 21 | 2 |
| Q12913 | Receptor-type tyrosine-protein phosphatase eta | 21 | 1 |
| O60449 | Lymphocyte antigen 75 | 0 | 1 |
| Q9H251 | Cadherin-23 | 0 | 1 |
| Q9H6A9 | Pecanex-like protein 3 | 0 | 1 |

\* From the search engine (Proteome Discoverer 2.4 software).

**Table 2.** Selected candidates for proximal membrane proteins around the SARS-CoV-2 spike protein in A549 cell surface.

| **Accession No.** | **Protein Name** | **Score\*** | **Peptide** |
| --- | --- | --- | --- |
| Q12860 | Contactin-1 | 231 | 5 |
| P01833 | Polymeric immunoglobulin receptor | 213 | 3 |
| Q86UN3 | Reticulon-4 receptor-like 2 | 77 | 4 |
| Q53RT3 | Retroviral-like aspartic protease 1 | 33 | 1 |
| P21796 | Voltage-dependent anion-selective channel protein 1 | 26 | 1 |
| Q6YHK3 | CD109 antigen | 22 | 1 |
| O00398 | Putative P2Y purinoceptor 10 | 0 | 1 |
| Q12864 | Cadherin-17 | 0 | 2 |

\* From the search engine (Proteome Discoverer 2.4 software).

Cadherin 17, which exhibited the highest score in Caco-2 cells, is a member of the cadherin superfamily that mediates Ca^2+^-dependent cell–cell adhesion without any connection to a cytoskeleton molecule. Cadherin 17 acts as a regulator of B cell development in knockout mice (38). Cadherin 17 can also influence the progression of certain cancers (39, 40). However, there is a paucity of studies on the relationship between Cadherin 17 and viral infection. Neither Prominin-1 (CD133) (41) nor vesicle-associated membrane protein-associated protein A (VAPA) (42) has been found to have a role in viral infection, including SARS-CoV-2 infection. Transferrin has been reported as a co-receptor for hepatitis C virus (43), but its relationship to SARS-CoV-2 infection has not yet been reported. In contrast, it has been found that DPP4 (44) plays a role in the pathogenesis of SARS-CoV-1 and/or SARS-CoV-2 infections (45). In A549 cells, CD44 (46), Contactin1 (47), Cadherin-1 (48), and Reticulon-4 receptor-like 2 (49) have not yet been reported to be involved in SARS-CoV-2 infection. In contrast, it has been reported that integrin families act as co-receptors for other viruses (50, 51), and it has subsequently been noted that these proteins exhibit a relationship with SARS-CoV-2 infection (52, 53). Interestingly, while carcinoembryonic antigen-related cell adhesion molecule (CEACAM) (54, 55) has been speculated to act as a primary receptor of mouse coronavirus (56), a recent network perturbation analysis suggested a very specific influence of this molecule on SARS-CoV-2 infection (57). Our findings, including those regarding CEACAM, and the observed concordance of our study results with those of previous studies indicate that potent candidate proteins can be obtained using the method described in this study.

The results of confocal microscopy experiment (Fig. 4) revealed that the candidate proteins co-localized with the spike protein. However, the co-localization was not identical. This could be attributed to the contribution of multiple factors (*e*.*g*., influence of the plasma membrane environment) other than the identified candidate proteins. It is unclear whether the extent of spike protein binding is directly associated with the possibility of SARS-CoV-2 infection.

A simple *in vitro* infection assay system was developed to elucidate whether the identified candidate molecules contribute to SARS-CoV-2 infection. VeroE6 or Caco-2 cells expressing TMPRSS2A (58) and others were considered for use as host cells in this study, but in order to observe the direct effect of the candidate molecule, we used stable transfectant HEK293T cells coexpressing ACE2 and candidate molecules. It was found that the proportion of the infected cells was approximately 1 to 5% of the total cells, and our finding was similar to the results reported in other studies (37, 59). In the infection assays using cells expressing each candidate molecule alone, almost no infection was observed (Fig. 6D), which indicated that CD133, CDH17, and VAPA were not the primary receptors for the SARS-CoV-2 spike protein. In contrast to this, the SARS-CoV-2 spike protein showed an increased binding capacity in coexpressing cells compared to that in cells expressing ACE2 alone. The number of infected cells with pSARS-CoV-2 also increased by 2-3 folds. CD133 expression showed a greater increase in this infection assay. Therefore, the results of this assay suggest that CD133, CDH17, and VAPA partially contribute to infection in ACE2 expressing cells. Although there was no statistically significant difference, there was an increase in SARS-CoV-2 spike protein binding to each candidate molecule-expressing P-ACE2, suggesting the role of the candidate molecules in assisting adhesion and/or entry of pSARS-CoV-2 virion into the host cell.

In conclusion, the candidate target proteins identified in this study were partially consistent with the results obtained from previous studies on SARS-CoV-2. Our method can therefore be considered an effective tool for identifying the host cell membrane proteins involved in viral infection. This strategy is likely to be applicable not only to SARS-CoV-2 but also to other viruses. This means that it can be used not only for known viruses but also for novel viruses that will emerge in the future. The results of this study will aid in the elucidation of entry mechanism of SARS-CoV-2 and other viruses, resulting in the development of novel therapeutic anti-viral agents in the future.

### Experimental Procedures

#### Cell culture

A549 human caucasian lung carcinoma and HEK293T human embryonic kidney cells (RIKEN CELL BANK, Saitama, Japan) were cultured in RPMI 1640 medium (Wako Chemicals, Miyazaki, Japan), supplemented with 5% fetal bovine serum (FBS; GIBCO bland, Thermo Fisher Scientific, MA), at 37°C under humidified air containing 5% CO_2_. Caco-2 cells (a gift from Dr. Sylvie Demignot, INSERM, Paris, France) were grown in Dulbecco’s modified Eagle’s medium (DMEM; Nacalai, Kyoto, Japan), supplemented with 4.5 g/L glucose, 10% FBS, and NEAA (Nacalai), at 37°C in a humidified atmosphere containing 5% CO_2_.

#### Preparation of Alexa Fluor 488-conjugated spike proteins

Recombinant SARS-CoV-2 spike protein (S1-RBD) was purchased from Sino Biological (40592-V05H; S1-RBD-mouse Fc, Beijing, China). Alexa Fluor 488-conjugated monovalent SARS-CoV-2 spike protein was prepared using 0.5 µg SARS-CoV-2 spike protein and the Zenon mouse IgG HRP labeling kit (Thermo Fisher Scientific) according to the manufacturer’s instruction. The negative control probe (cAb-488) was prepared using non-specific mouse IgG fraction included in the labeling kit.

#### Immunocytochemistry

For immunocytochemistry, Caco-2 and A549 cells were grown on 35 mm glass bottom dishes (Matsunami Glass, Osaka, Japan). The cultured cells were fixed with or without 4% paraformaldehyde, washed thrice with PBS, and then stained with the Alexa Fluor 488-conjugated monovalent SARS-CoV-2 spike protein at room temperature for 30 min. In addition, the cells were stained with the SARS-CoV-2 spike protein (40592-V05H; 2.5 µg/ml 2% BSA-PBS), followed by the secondary antibody anti-mouse IgG Alexa Fluor 488 (A-11001; Thermo Fisher Scientific; 5 µg/ml 2% BSA-PBS). To confirm the expression of ACE2 in Caco-2 and A549 cells, the cultured cells were fixed with 4% paraformaldehyde, washed thrice with PBS, and then stained with an anti-ACE2 antibody (PAB886Hu01; CLOUD-CLONE, Wuhan, China; 5 µg/ml 2% BSA-PBS) at room temperature for 30 min, followed by incubation with anti-rabbit IgG Alexa Fluor 568 (ab175471; Abcam, Cambridge, UK; 10 µg/ml 2% BSA-PBS) at room temperature for 30 min. The stained samples were observed with a fluorescent microscope (BZ-700, Keyence, Osaka, Japan). Raw images, including the differential interference contrast image, were captured under identical settings, in the case of same experiments, and then exported as TIFF files.

#### Western blotting

Samples solubilized with a reducing SDS sample buffer or the eluted samples from purified and enriched resins described above were subjected to SDS-PAGE using 10% gels. After electrophoresis, the gels were blotted onto an Immobilon®-P PVDF Membrane (Millipore, MA), followed by blocking with a 5% skim milk solution. The membranes were then incubated with the following primary antibodies: anti-ACE2 antibody (PAB886Hu01; CLOUD-CLONE; 0.5 µg/ml 5% skim milk solution) and HRP-conjugated anti-fluorescein antibody (SouthernBiotech, AL; 0.5 µg/ml 5% skim milk solution) described in a previous study (31); the incubations were carried out at 4°C overnight or at room temperature for 1 h, respectively. In the case of ACE2 antibody, membranes were incubated with a secondary antibody, HRP-conjugated anti-rabbit IgG (0.4 µg/ml 5% skim milk solution), at room temperature for 1 h. Membranes were developed with the Immobilon Western Chemiluminescent HRP Substrate (Millipore) and analyzed using a ChemiDoc MP Image analyzer (BioRad, CA). The molecular weight marker was a Pre-stained Protein Markers, Broad Range (Nacalai). For loading controls, the PVDF membrane after exposure was stained with Coomassie Brilliant Blue solution.

#### EMARS reaction

The EMARS reaction and the detection of EMARS products were performed as described previously (20). Briefly, Caco-2 and A549 cells were cultured in 10 cm plastic culture dishes (TPP, Trasadingen, Switzerland), until they were approximately 80-90% confluent, and then washed once with PBS at room temperature and subsequently treated with HRP-conjugated Cholera Toxin B Subunit B (CTxB-HRP; LIST Biological Lab, CA) or SARS-CoV-2 spike protein (0.5 µg/ml 2% BSA-PBS) at room temperature for 30 min. After washing thrice with PBS, the cells were treated with HRP-conjugated anti-mouse IgG (W402B; Promega, WI; 0.25 µg/ml 2% BSA-PBS) at room temperature for 30 min. After gently washing five times with PBS, the treated cells were then incubated with 0.05 mM fluorescein-conjugated tyramide (FT) (21), containing 0.0075% H_2_O_2_ in PBS, at room temperature for 10 min in the dark. The cell suspension was homogenized through a 22 G syringe needle to rupture the plasma membranes, and the samples were centrifuged at 20,000 *g* for 15 min to precipitate the plasma membrane fractions. After solubilization using the NP-40 lysis buffer (20 mM Tris-HCl (pH 7.4), 150 mM NaCl, 5 mM EDTA, 1% NP-40, 1% glycerol), the samples were subjected to SDS-PAGE as described above.

#### Purification and enrichment of EMARS products

Following the EMARS reaction, the precipitated cell membrane pellet was mixed with chloroform and methanol in a 2:1 volume ratio. Deionized water was then added, and gentle agitation was done. All the solvent was removed, and the resulting pellets were washed thrice with 40% methanol to completely remove excess FT. To remove any residual solution completely, the specimens were evaporated and solubilized with 100 µL of 50 mM Tris-HCl (pH 7.4), containing 1% SDS, at 95°C for 5 min. The soluble material was transferred into a new tube and then diluted with 400 µL NP-40 lysis buffer. Then, 20 µL of the prepared anti-fluorescein antibody Sepharose, which was produced by the conjugation reaction between anti-fluorescein antibody (SouthernBiotech) and NHS-activated Sepharose 4 Fast Flow (GE Healthcare Life Sciences, MA), was added to the sample, which was mixed with rotation using rotator at 4°C overnight. After washing resins with the NP-40 lysis buffer five times and 0.5 M NaCl-PBS two times, a 1% SDS solution, containing MPEX PTS reagent (GL Sciences, Tokyo, Japan), was added to resins for MS analysis. The samples were then heated at 95°C for 5 min to elute the fluorescein-labeled molecules from resins.

#### Proteomic analysis of EMARS products using mass spectrometry analysis

Proteomic analysis was performed using nano-liquid chromatography-electrospray ionization mass spectrometry (nano LC-ESI-MS/MS). The eluted samples described above were treated with a final 10 mM concentration of DTT (Wako Chemical) at 50°C for 30 min and then treated with 50 mM iodoacetamide (Wako Chemical) in 50 mM ammonium carbonate buffer at 37°C for 1 h. To remove SDS, a 5% v/v SDS-eliminant (ATTO, Tokyo, Japan) was directly added to the samples, and the samples were incubated at 4°C for 1 h. After centrifugation, the supernatant was transferred to a new tube and then digested with 2 µg trypsin (Trypsin Gold MS grade; Promega) at 37°C overnight. For peptide purification, the digested sample was applied to a C18-StageTip (GL Sciences) according to the manufacturer’s instructions. The sample was solubilized in 10 µL of a 2% acetonitrile (ACN)/0.1% formic acid/H2O solution. The prepared samples were then injected into Ultimate 3000 RSLCnano (Thermo Fisher Scientific). Mass analysis was performed using a LTQ Orbitrap XL mass spectrometer equipped with a nano-ESI source (Thermo Fisher Scientific). A NIKKYO nano HPLC capillary column (3 μm C18, 75 μm I.D. × 120 mm; Nikkyo Technos, Tokyo, Japan) was used, along with a C18 PepMap100 column (300 μm I.D. × 5 mm; Thermo Fisher Scientific). The peptides were eluted from the column using a 4–35% acetonitrile gradient for over 97 min. The eluted peptides were directly electrosprayed into the spectrometer using a precursor scan and data-dependent MS/MS scan. The mass spectrometry system parameters and search parameters used in this study are summarized in Table S6. For each raw data file recorded by the mass spectrometer, peak lists were generated using Proteome Discoverer ver. 2.2 or 2.4 (Thermo Fisher Scientific). Peak lists, generated by Proteome Discoverer, were searched against the Swiss-Prot database (taxonomy: *Homo sapiens*) using the Mascot search engine. The false discovery rate (FDR) was calculated using the peptide and protein decoy databases in Proteome Discoverer. The FDR was strict 1% and relaxed 5% at the peptide and protein level. The MS data sets (including RAW and result files) were posted to the Japan ProteOme STandard (jPOST) Repository/Database (https://jpostdb.org/).

The EMARS products from A549 and Caco-2 cells were subjected to MS analysis in duplicates, and then suitable candidate proteins were filtered based on the following exclusion criteria: (i) keratin, immunoglobulin, histone, albumin, trypsin, and actin (ii) proteins detected in negative control samples (spike [-] sample).

#### Confocal Microscopy for Caco-2 cells

Caco-2 cells were seeded and cultured onto 8-well format slides for confocal analysis (SARSTEDT). The slides were then washed with PBS once and treated with the SARS-CoV-2 spike protein (0.5 µg/ml 2% BSA-PBS) at room temperature for 30 min. After fixing with 4% paraformaldehyde at room temperature for 15 min, the specimens were gently washed with PBS, followed by addition of the following primary antibodies and incubation at room temperature for 30 min: anti-ACE2 antibody (PAB886Hu01; CLOUD-CLONE; 5 µg/ml 2% BSA-PBS), anti-DPP4 antibody (10940-1-AP; PROTEINTECH, Tokyo, Japan; 1 µg/ml 2% BSA-PBS), anti-CD133 antibody (18470-1-AP; PROTEINTECH; 1.5 µg/ml 2% BSA-PBS), or anti-Cadherin 17 antibody (CSB-PA006407; CUSABIO, Wuhan, China; 1.5 µg/ml 2% BSA-PBS). After washing thrice with PBS, the samples were treated with the secondary antibodies anti-mouse IgG Alexa Fluor 488 (5 µg/ml 2% BSA-PBS) and anti-rabbit IgG Alexa Fluor 568 (10 µg/ml 2% BSA-PBS) at room temperature for 30 min. The stained slides were fixed with ProLong™ Glass Antifade Mountant (Thermo Fisher Scientific). The samples were observed with an LSM 710 Laser Scanning Confocal Microscope (Carl Zeiss, Oberkochen、Germany) mounted on an AxioImager Z2 equipped with a Diode laser unit (405 nm/30 mW), Argon laser unit (458, 488, 514 nm/25 mW), He-Ne laser unit (543 nm/1 mW) and He-Ne laser unit (633 nm/5 mW). The objective lenses were EC-PLAN NEOFLUAR 5x/0.16 and APOCHROMAT 20×/0.8 (Carl Zeiss). Image acquisition and analysis were carried out with ZEN 2011 software (Carl Zeiss). Raw images, including the differential interference contrast image, were captured under the identical settings in the case of same experiments and then exported as TIFF files.

#### Transmission electron microscopy (TEM) for Caco-2 cells

Caco-2 cells were grown on 35 mm glass bottom dishes (Matsunami Glass) and pre-fixed with 4% paraformaldehyde in PBS for 20 min. Then, the cells were stained with spike protein (5 μg/ml) in 1% BSA-PBS for 30 min, followed by incubation with the 20 nm gold particle-conjugated anti-mouse IgG antibody (BBI international, Crumlin, UK; EM. GAT10: 1:100) for 20 min. The samples were fixed again and then stained with an anti-CD133 antibody (18470-1-AP; 1.5 µg/ml), anti-DPP4 antibody (10940-1-AP; 1 µg/ml), anti-Cadherin 17 antibody (CSB-PA006407; 1.5 µg/ml), or anti-VAPA antibody (15275-1-AP; PROTEINTECH; 4.2 µg/ml) for 30 min, followed by incubation with the 10 nm gold particle-conjugated anti-rabbit IgG antibody (BBI international; EM.GAT5: 1:100) for 30 min. The samples were then fixed with 2% glutaraldehyde-PBS at 4°C overnight, dehydrated using a graded series of ethanol (30%–100%), and embedded in epoxy resin. Next, the samples were cut with an Ultramicrotome (EM UC7, Leica, Hessen, Germany), stained with uranyl acetate, and observed with a transmission electron microscope (Talos L120, Thermo Fisher Scientific).

#### Preparation of HEK293T transfectant cells

HEK293T cells were cultured in a 6-well dish (approximately 80% confluent), and 1 μg of pCMV3-human ACE2 (HG10108-UT; Sino Biological) was transfected using 5 μl TransIT^®^-2020 Reagent (Mirus Bio, WI). After 3 days, 150 μg/ml hygromycin was added, and the cells were cultured for approximately 2 weeks. A few colonies were obtained and transferred into a new 12-well culture plate. One of these cell colonies was used as P-ACE2.

Human ACE2 ORF cassette was obtained via enzymatic digestion of the pCMV3-ACE2 vector with KpnI and XbaI. The ACE2 ORF cassette was inserted into pENTR1A no ccDB (w48-1) (addgene, MA). It was recombined into pLenti CMV/TO Puro DEST (addgene) using the Gateway LR Clonase Enzyme mix (Thermo Scientific). This pLenti-ACE2 vector, pMD2.G (addgene), and psPAX2 (addgene) were transfected together into HEK293T cells using the TransIT^®^-2020 Reagent. After 72 h of culture, the culture supernatant containing lentivirus was harvested and then passed through a 0.45 μm filter (Sartorius, Niedersachsen、Germany) to remove cell debris.

The HEK293T cells cultured in 6-well dish (approximately 50% confluent) were subsequently treated with 1 ml of this virus solution for 3 days. The cultured cells were then treated with 2 μg/ml puromycin and cultured for another 2 weeks. The cells selected by puromycin treatment were designated as L-ACE2 cells.

The ORFs of candidate proteins, CD133 and VAPA, were obtained via PCR cloning using the human cDNA library from HepG2 cells. The ORFs of the negative control protein GPC3 were also obtained by PCR cloning using the human cDNA library from HepG2 cells. For PCR cloning, we used KOD plus DNA polymerase (TOYOBO, Osaka, Japan) and DNA primers as follows: CD133-5′-kpnI, acaggtaccatggccctcgtactcggctc; CD133-3′-XhoI, tgtctcgagtcaatgttgtgatgggcttg; VAPA-5′-KpnI, acaggtaccatggcgtccgcctcaggggc; VAPA-3′-XhoI, tgtctcgagctacaagatgaatttcccta; GPC3-5′-KpnI, acaggtaccatggccgggaccgtgcgcac; GPC3-3′-XhoI, tgtctcgagtcagtgcaccaggaagaaga. The ORFs of the candidate protein CDH17 were obtained from pCMV3-human CDH17 (HG11360-UT; Sino Biological) by KpnI and XbaI digestion. These ORF cassettes were inserted into pENTR1A no ccDB (w48-1), followed by recombination into pLenti CMV/TO Puro DEST for the construction of lentivirus vector for candidate protein expression. Lentivirus production for the expression of these candidate proteins was performed in the same way as for ACE2 expression lentivirus. For the preparation of coexpressing cells (P-ACE2-CD133, P-ACE2-CDH17, P-ACE2-VAPA, and P-ACE2-GPC3), the obtained lentivirus solution was treated with P-ACE2 cells, followed by selection using 2 μg/ml puromycin.

Each cell was stained with an anti-ACE2 antibody (PAB886Hu01; 5 µg/ml 2% BSA-PBS), anti-CD133 antibody (18470-1-AP; 1.5 µg/ml), anti-Cadherin 17 antibody (CSB-PA006407; 1.5 µg/ml), and anti-VAPA antibody (15275-1-AP; 4.2 µg/ml), followed by incubation with Alexa Fluor 488-conjugated anti-rabbit IgG (ab150077; Abcam; 2.5 µg/ml). After washing with PBS, the treated cells were analyzed, using a BD FACS Canto II flow cytometer, to determine the expression of each molecule (Becton Dickinson, NJ). Mean fluorescence intensity (MFI) was used for the comparison of expression levels.

#### In vitro infection assay

SARS-CoV-2 pseudovirus (pSARS-CoV-2), which has a gene cassette for expression of Green Fluorescent Protein (GFP), was prepared using the pPACK-SPIKE™ SARS-CoV-2 “S” Pseudotype Lentivector Packaging Mix (Wuhan-Hu-1; CVD19-500A-1; SBI) according to manufacturer’s instructions. Briefly, HEK293T cells (approximately 50% confluent) were cotransfected with the pPACK vector solution and pLenti CMV/TO eGFP Puro (w159-1) (addgene) using the TransIT^®^-2020 Reagent as described above. After 72 h of culture, the culture supernatant containing pSARS-CoV-2 was harvested and then passed through a 0.45 μm filter to remove cell debris. The Lenti-X™ Concentrator (Clontech) was added (1 volume of Lenti-X Concentrator with 3 volumes of clarified supernatant), followed by incubation at 4 °C overnight. After centrifugation at 1,500 *g* for 45 min at 4 °C, the supernatant was removed, and pSARS-CoV-2 pellet was gently resuspended in 500 μl of RPMI 1640 containing 5% FBS.

The host cells (P-ACE2-CD133 cell etc.; 1.5 × 10^5^ cells/well) and 50 μl of supernatant containing pSARS-CoV-2 prepared above were mixed and transferred into a 12-well dish. After 60 h of culture, the infected cells expressing GFP were subjected to BD FACS Canto II for counting the GFP-positive cells. The GFP-positive pSARS-CoV-2 cells were considered as infected cells if they had a GFP fluorescence intensity of 10^3^ or higher. To reduce the possibility of error, a total of 100,000 cells were counted. In addition, the infected cells were simultaneously observed with a EVOS FLoid® Cell Imaging Station fluorescence microscope (Thermo Fisher Scientific).

### Data processing

Statistical analyses were performed using the R software (The R Foundation for Statistical Computing, Austria) and EZR (Saitama Medical Center, Jichi Medical University, Japan) (60), a graphical user interface for R. Statistical significance tests for the experiments of spike protein binding and *in vitro* infection assay were performed using the Dunnett’s test in R. A p-value ≤ 0.05 was considered statistically significant.

## Data availability

All data are contained within the article.

## Acknowledgements

We thank Ms. Amimoto of the Natural Science Center for Basic Research and Development (N-BARD), Hiroshima University, for MS technical assistance and the Saitama Medical University Biomedical Research Center for providing general technical assistance.

## SUPPORTING INFORMATION

Table S1 Raw data of MS analysis for Caco-2 cells; first MS analysis

Table S2 Raw data of MS analysis for Caco-2 cells; second MS analysis

Table S3 Raw data of MS analysis for A549 cells; first MS analysis

Table S4 Raw data of MS analysis for A549 cells; second MS analysis

Table S5 Identified membrane proteins involved in SARS-CoV-2 infection

Table S6 Mass spectrometry system parameters and search parameters used in this study

**Fig S1.**
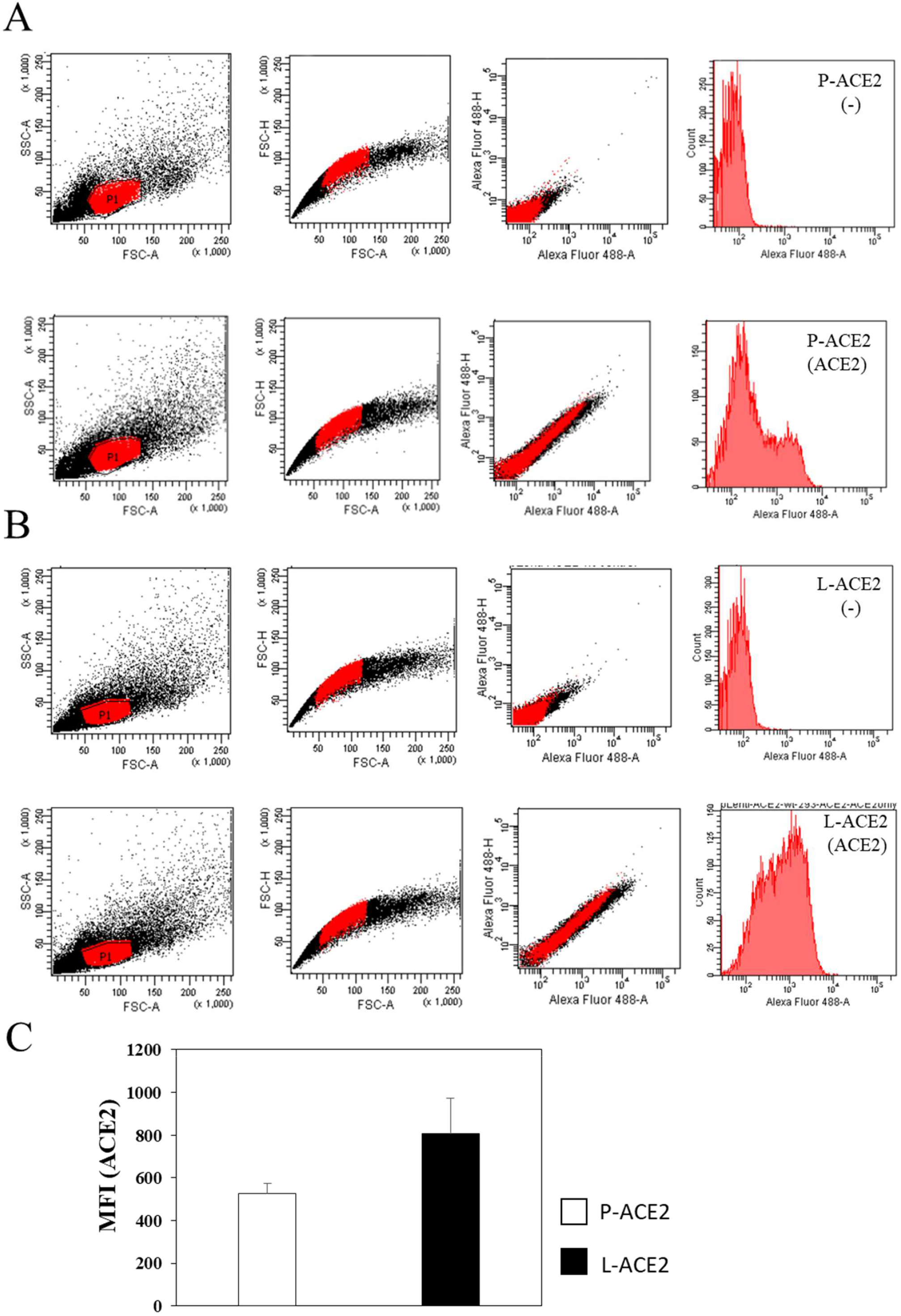
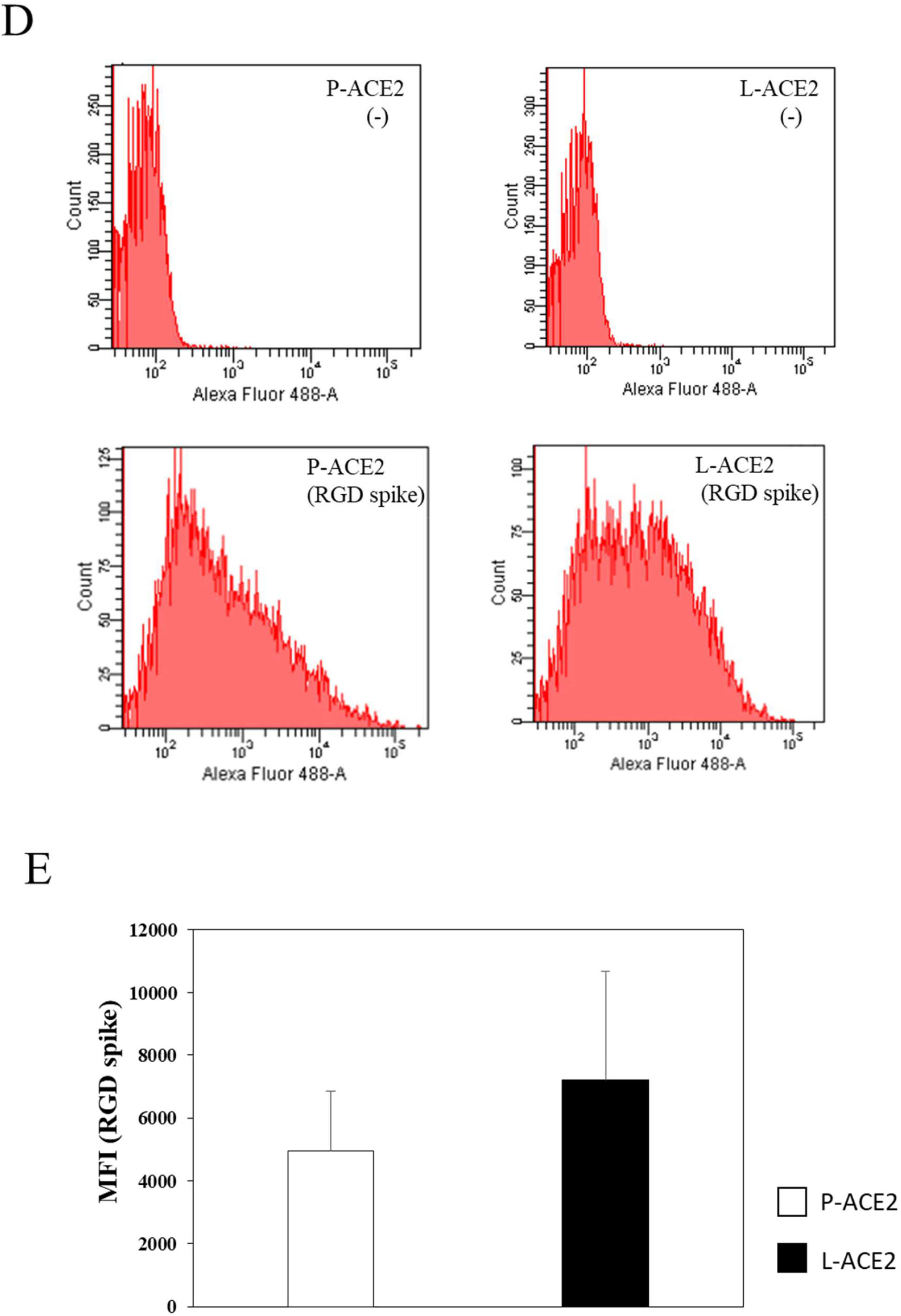
Characterization of ACE2-expressing HEK293T cells. (*A, B*) Representative data from flow cytometric analysis of P-ACE2 (*A*) and L-ACE2 (*B*) cells. P-ACE2 and L-ACE2 cells were treated with an anti-ACE2 antibody, followed by incubation with Alexa Fluor 488-conjugated anti-rabbit IgG (P-ACE2 (ACE2) and L-ACE2 (ACE2); lower panel), or without anti-ACE2 antibody (P-ACE2 (-) and L-ACE2 (-); upper panel). The samples were subsequently subjected to flow cytometric analysis (Forward Scatter (FSC) vs. Side Scatter (SSC) plots, Alexa Fluor 488 plots, and Histogram of cell count vs. fluorescein intensity). Three independent experiments were performed. (*C*) Comparison of the mean fluorescence intensity (MFI) between P-and L-ACE2 cells. The MFI (ACE2) in L-ACE2 cells with anti-ACE2 antibody staining (closed bar) was higher than that in P-ACE2 cells (open bar). (*D*) Representative data from flow cytometric analysis of the spike protein-treated P-ACE2 and L-ACE2 cells. The cells were treated with SARS-CoV-2 spike protein, followed by incubation with Alexa Fluor 488-conjugated mouse IgG second antibody (P-ACE2 (RGD spike) and L-ACE2 (RGD spike); lower panel), or without spike protein (P-ACE2 (-) and L-ACE2 (-); upper panel). The cells were subsequently subjected to flow cytometric analysis. Three independent experiments were performed. (*E*) The MFI (RGD spike) in L-ACE2 cells (closed bar) with spike protein staining was higher than that in P-ACE2 cells (open bar).

**Fig. S2.**
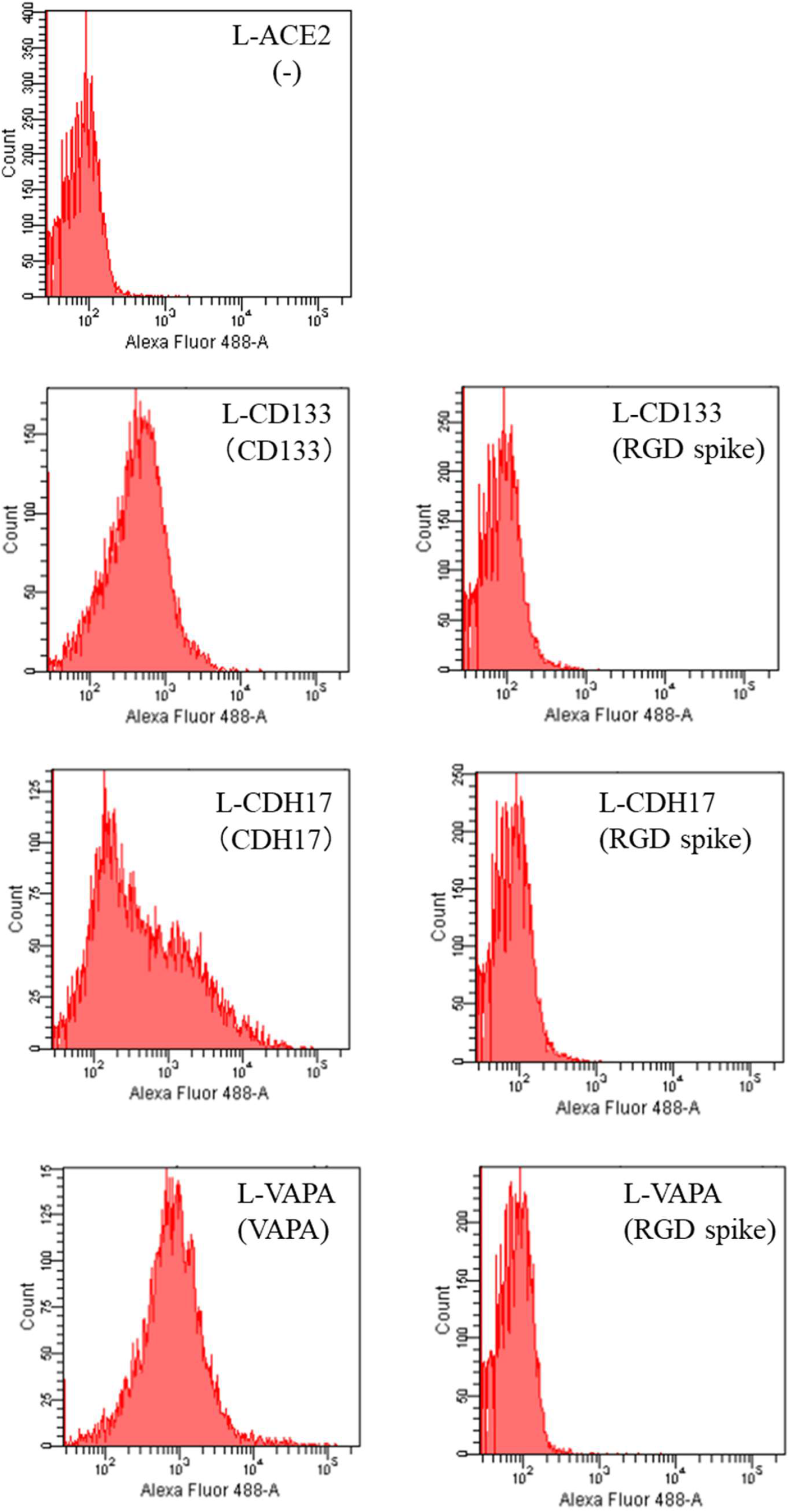
Characterization of the single candidate molecule-expressing HEK293T cells. Representative data from flow cytometric analysis of L-CD133, L-CDH17, and L-VAPA cells. Firstly, these cells were treated with antibodies against each candidate molecule, followed by incubation with Alexa Fluor 488-conjugated anti-rabbit IgG (L-CD133 (CD133), L-CDH17 (CDH17), and L-VAPA (VAPA); left panel). The cells were also treated with spike protein, followed by Alexa Fluor 488-conjugated anti-mouse IgG (L-CD133 (RGD spike), L-CDH17 (RGD spike), and L-VAPA (RGD spike); right panel). Two independent experiments were performed.

**Fig. S3.**
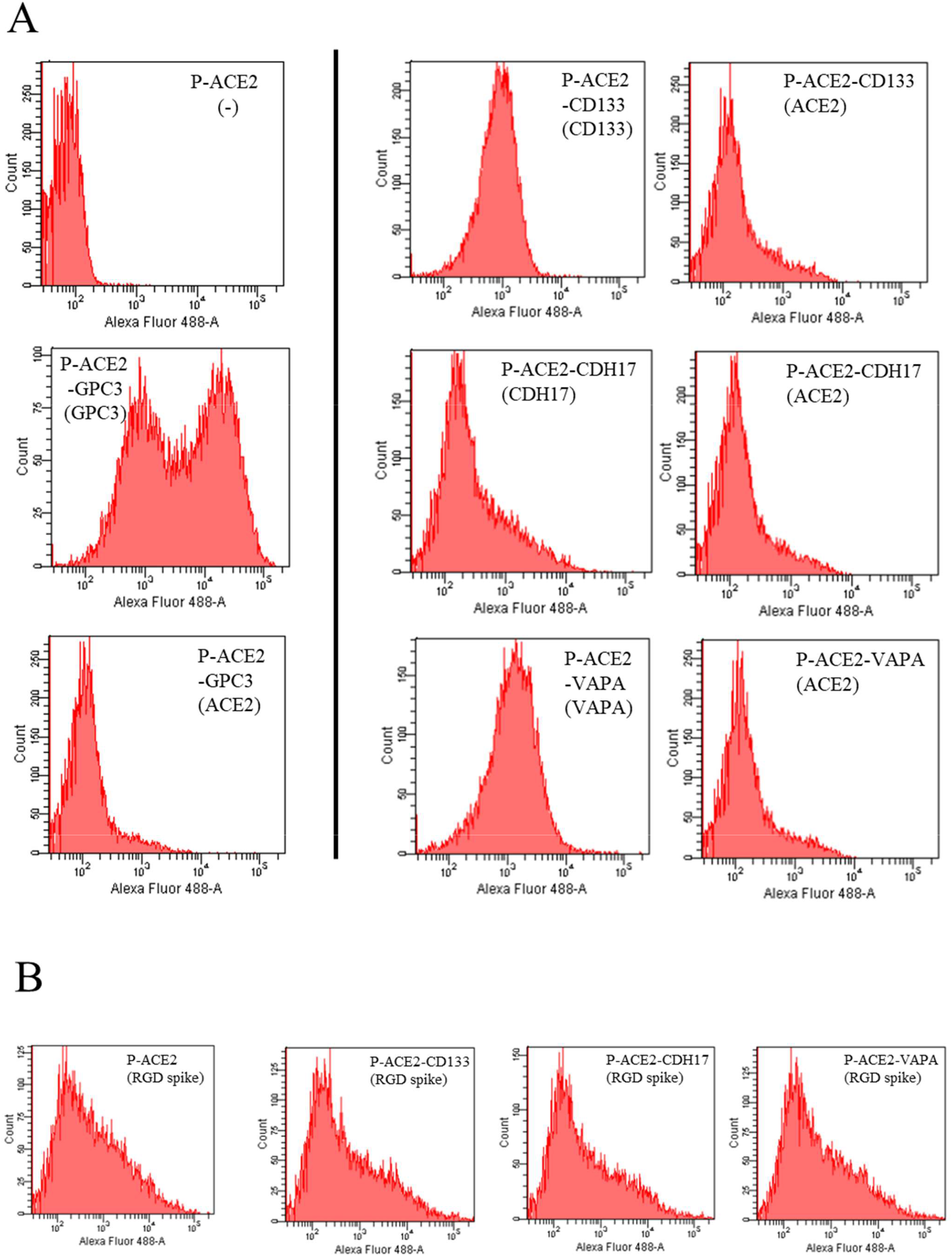
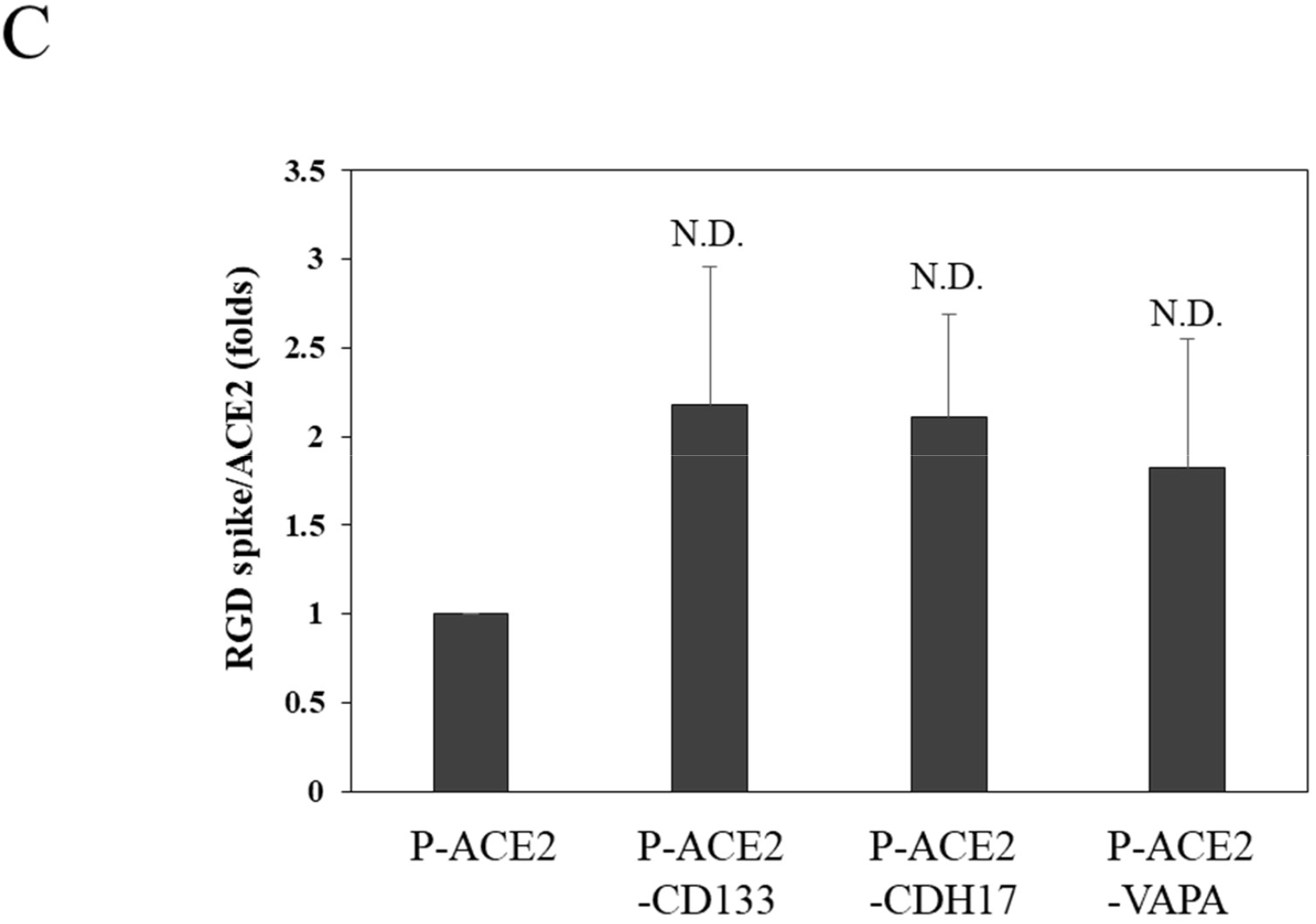
Characterization of the ACE2- and candidate molecule-coexpressing HEK293T cells. (*A, B*) Representative data from flow cytometric analysis of ACE2- and candidate molecule-coexpressing cells. (*A*) The cells were treated with an anti-ACE2 antibody (P-ACE2-CD133 (ACE2), P-ACE2-CDH17 (ACE2), P-ACE2-VAPA (ACE2), and P-ACE2-GPC3 (ACE2)), first with antibody recognizing each candidate molecule (P-ACE2-CD133 (CD133), P-ACE2-CDH17 (CDH17), P-ACE2-VAPA (VAPA) and P-ACE2-GPC3 (GPC3)), and then with Alexa Fluor 488-conjugated anti-rabbit IgG. Three independent experiments were performed. (*B*) The cells were treated with SARS-CoV-2 spike protein (RGD), followed by Alexa Fluor 488-conjugated mouse IgG second antibody (P-ACE2 (RGD spike), P-ACE2-CD133 (RGD spike), P-ACE2-CDH17 (RGD spike), and P-ACE2-VAPA (RGD spike)). Three independent experiments were performed. (*C*) The relative MFI values (RGD spike/ACE2) of P-ACE2-CD133, P-ACE2-CDH17, and P-ACE2-VAPA cells were higher than those of P-ACE2 cells, but no significant difference (N.D.) was observed.

